# Extant hybrids of RNA viruses and viroid-like elements

**DOI:** 10.1101/2022.08.21.504695

**Authors:** Marco Forgia, Beatriz Navarro, Stefania Daghino, Amelia Cervera, Andreas Gisel, Silvia Perotto, Dilzara N. Aghayeva, Mary Funmilayo Akinyuwa, Emanuela Gobbi, Ivan N. Zheludev, Robert C. Edgar, Rayan Chikhi, Massimo Turina, Artem Babaian, Francesco Di Serio, Marcos de la Peña

## Abstract

Earth’s life may have originated as self-replicating RNA. Some of the simplest current RNA replicators are RNA viruses, defined by linear RNA genomes encoding an RNA-dependent RNA polymerase (RdRP), and subviral agents with single-stranded, circular RNA genomes, such as viroids encoding paired self-cleaving ribozymes. Amongst a massive expansion of candidate viroid and viroid-like elements, we report that fungal pathogens, ambiviruses, are viroid-like elements which undergo rolling circle replication and encode their own viral RdRP, thus they are a distinct hybrid infectious agent. These findings point to a deep evolutionary history between modern RNA viruses and sub-viral elements and offer new perspectives on the evolution of primordial infectious agents, and RNA life.

**One-Sentence Summary:** Novel infectious agents resembling self-cleaving viroid-like RNAs whilst encoding a viral RNA-dependent RNA polymerase.

---

Serving dual functions as encoding for genetic information, and as a biological catalyst, RNA has been proposed to predate DNA and protein at the origin of life in an “RNA World” (*1, 2*). Further, extant sub-viral and viral agents have been suggested to be “living fossils” and can help illuminate the molecular biology of Earth’s first life (*3-5*).

Viruses having RNA genomes (realm *Riboviria*) are infectious agents defined by a linear RNA genome encoding one of their hallmark replication polymerases (replicases), either an RNA dependent RNA polymerase (RdRP) for RNA viruses, or a Reverse Transcriptase (RT) for retroviruses. These replicases, and thus *Riboviria*, are monophyletic in nature as based on alignment of the conserved palm domain of RdRP and RT(*6*).

Viroids and viroid-like entities (such as the human satellite virus, Hepatitis Delta Virus), are sub-viral RNA infectious agents of plants and animals (Fig 1A). These sub-viral agents are typified by single-stranded circular RNA (circRNA) genomes which form extensive secondary structures, typically rod or branched-rod folding, and most encode hallmark paired ribozymes, one in each strand polarity, necessary for their rolling-circle replication. So far, the self-cleaving RNA ribozymes reported in infectious circRNA (*7, 8*) are hammerhead (HHRz), hairpin (HPRz) and hepatitis Delta (DVRz) ribozymes (*9, 10*). In contrast to RNA viruses, infectious circRNAs replicate by RNA polymerases encoded *in trans* by the host or helper viruses. Additionally, while viroids are non-coding RNAs (8), Delta and Delta-like viruses have been assigned the realm *Ribozyviria*, and encode for the Delta Antigen proteins (*11*). Overall, the simple genomes and small ribozymes of infectious circRNAs are why these elements are regarded as molecular fossils of the prebiotic world (*3*).

**Fig. 1:**
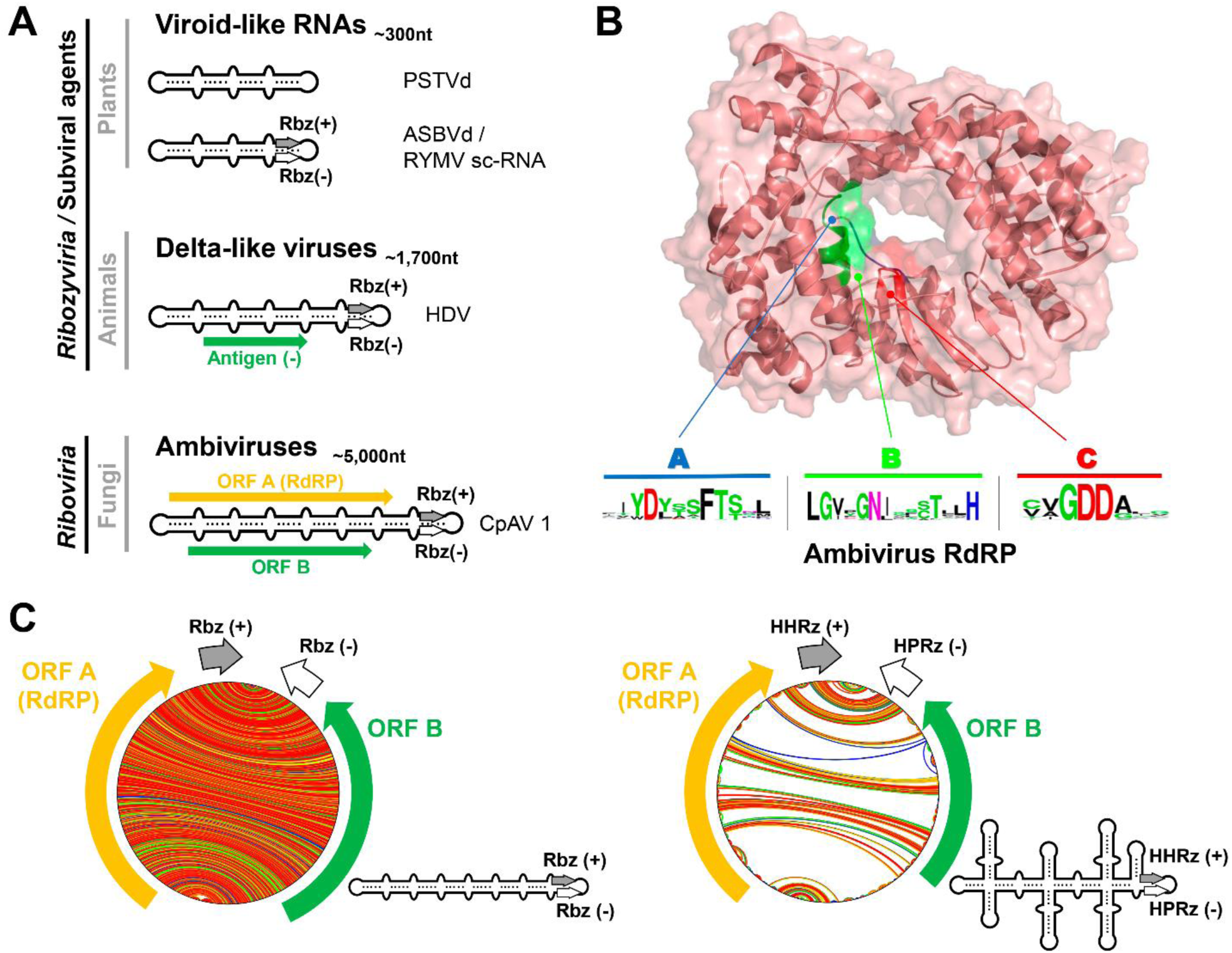
Sub-viral circRNA and ambivirus genome organizations and ambivirus RNA-dependent RNA polymerase. (**A**) Schematic genome organization of representative sub-viral agents: non-coding viroids and viroid-like satellite RNAs; and protein-coding Delta-like viruses (*Ribozyviria)* and ambiviruses. (**B**) (AlphaFold2 predicted structure of the representative Armillaria borealis ambivirus 2 (Accession MW4238010.1) ORF-A shows a classic polymerase palm fold structure and strong sequence conservation of catalytic residues in motif A, B, and C is seen across all ambiviruses. (**C**) Circular plots of the predicted RNA secondary structures of two representative ambiviruses. On the left, an example of the characteristic rod-like structure predicted for most ambiviruses. On the right, an example of a highly branched architecture predicted for some ambivirus RNA genomes, usually carrying a mixed pair of HHRz/HPRz motifs.

Ambiviruses are a recently characterized and widespread group of fungal single-stranded RNA infectious agents with unusual genomic features (*12, 13*) (Fig. 1A). An ambivirus (∼5 kb genome) encodes for two conserved open read frames (ORFs) A and B, one on each strand – a so-called bicistronic, ambisense genome. ORF-A has a remote similarity to RdRP leading to these agents being reported as RNA viruses. However, Northern blotting, RT-PCR, and *de novo* transcript assembly indicated contiguity between 3’ and 5’ terminal ends, inconsistent with the assumed typical linear RNA virus genome (*12, 13*).

Here we report that ambiviruses have circular RNA genomes, encoding paired self-cleaving ribozymes, enabling a viroid-like rolling circle mechanism of replication, however, we further show that ORF-A is indeed a functional RdRP like that of a classical RNA virus. Thus, ambiviruses appear to have arisen from the hybridization of a viroid-like genomic backbone with the hallmark RNA virus gene RdRP, and as such are a distinct class of infectious agent. This finding bridges the RNA virospheres of viruses and viroids, posing deep questions regarding their origins and (co-)evolution.

Starting with the hint that ambiviruses may have non-standard linear RNA genomes (Supplementary text and fig. S1), we analyzed published ambiviruses for sequence homology to structured RNA covariance models with INFERNAL (*14*). This uncovered head-to-tail oriented, ambisense HHRz or HPRz motifs in 28/30 (93%) of the GenBank sequences, reminiscent of subviral circRNAs (table S1). To expand the known sequence diversity of ribozyme-bearing subviral agents and ambiviruses, we adapted the *Serratus* ultra-high throughput computing architecture (*15*) to search for ribozymes using INFERNAL across 198,194 raw metagenome/metatranscriptomes freely-available in the Sequence Read Archive (*16, 17*). Combining the top 5,000 ribozyme-hit libraries with our previous RNA virus assemblage and the Transcriptome Shotgun Assembly (TSA), we created the “RNA Deep Virome Assemblage” (RDVA) of 58,557 libraries. From the 12.5 billion assembled contigs, 34 million contained 5’-3’ k-mer overlaps suggestive of a circular molecule. Filtering the potential circular contigs further for those encoding two paired-ambisense ribozymes resulted in a discovery set of 32,393 putative viroid and viroid-like elements, which clustered into 20,364 novel species-like operational taxonomic units (sOTU) at 90% nucleotide sequence identity. These datasets included up to 863 unique RdRP-encoding ambiviruses, which clustered into 378 sOTU at 90% palmprint identity (*18*).

The initially reported homology between ambivirus ORF-A and RdRP was marginal owing to their deep divergence (*12, 13*). Advances in *in silico* structural prediction have extended the capacity to assess for deep protein homology from sequences alone (*19*). AlphaFold2 prediction of ORF-A strongly supports (pLDDT confidence up to 93.5 for several models) that this protein takes on the classic right-handed palm domain architecture, with highest similarity to negarnavirus RdRPs (Influenza A, Z-score >19, RDSM 3.8A) (fig. S2). Critically, the expanded multiple sequence alignment shows the strong sequence conservation across the essential polymerase catalytic motifs A, B, and C (*18*) (Fig. 1B). Altogether this lends strong support that the ambivirus ORF-A encodes for a catalytically active RdRP.

Further screening of public databases (see Material and Methods) extended the number of ambiviral sOTUs up to 439 species. Prediction of the secondary structures of minimal free energy for the ambiviral potential circular genomes revealed than the majority (90%) of these RNAs adopt a perfect rod or quasi-rod like structure of very high stability, akin to most plant viroids and Delta-like viruses. However, about 10% of the ambivirus genomes adopt a highly branched RNA conformation, still stable and somehow keeping a main rod-like architecture (Fig. 1C), which suggests that extensive genomic base-pairing structural constraints are preserved across ambiviruses.

Reconstruction of the ambivirus RdRP phylogeny shows a complex natural history of the ambivirus RdRP and their ribozymes. Four out of the ten known self-cleaving ribozymes (*20*) were found in ambivirus genomes thereby suggesting that multiple recombination and/or horizontal transfer events of ribozymes have occurred (Fig. 2). Intriguingly, there is also extensive mixture of two different ribozymes on a single genome. Similar ribozyme mixtures are observed in the 5% of our discovery set of small viroid-like genomes, supporting that different ribozyme classes may coexist within a genome.

**Fig. 2:**
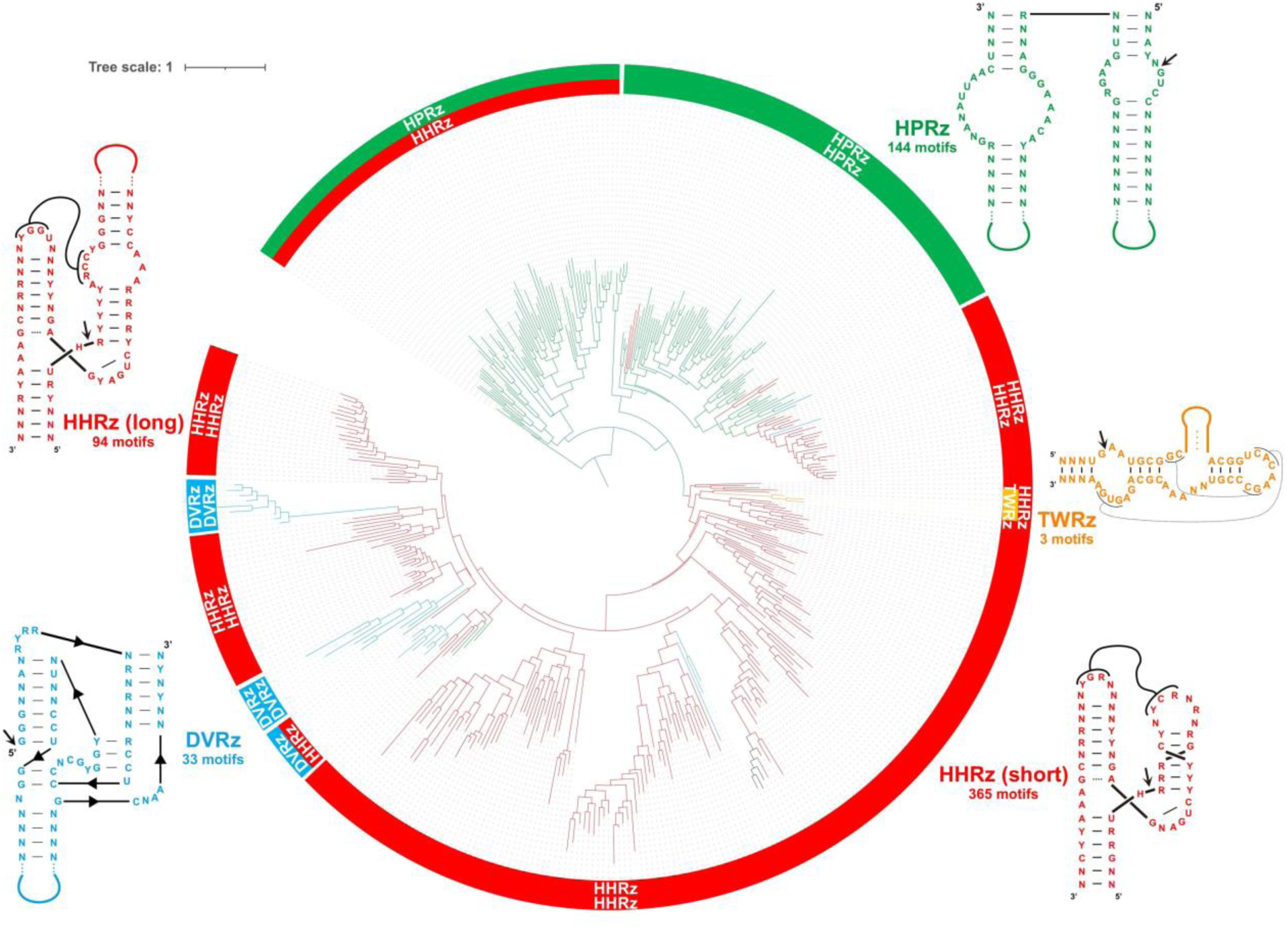
Ambivirus RNA-dependent RNA polymerase phylogeny. Maximum-likelihood phylogenetic tree of the RNA-dependent RNA polymerase palmprints from the 439 distinct species-like operational taxonomic units (sOTUs). There are six major clades of ambiviruses showing diverse ribozyme usage. Up to 60% sOTUs use paired type III HHRz similar to the motifs from Epsilonviruses (*15, 21*) or plant viroids (*3*), whereas ∼30% carry either two HPRz or a HHRz/HPRz, 5% carry the DVRz (either paired or a mix with HHRz) characteristic of animal deltaviruses, and three sOTUs contain a mix of the TWR (P1 architecture) (*22*) and the HHRz motifs. Consensus ribozyme structures (weighted nucleotide conservation threshold of 70%) of the hammerhead (HHRz, in red, two length variants), Delta virus (DVRz, in blue), hairpin (HPRz, in green) and twister ribozymes (TWR, in orange) present in ambiviruses are shown.

To validate this *in silico* expansion of ambiviruses, we sought to molecularly evaluate key properties from the predicted sequences. Self-cleavage capacity of ambivirus ribozymes was confirmed by *in vitro* transcription of their cDNAs from *Cryphonectria parasitica* ambivirus 1 (CpAV1) (two HHRzs), Tulasnella ambivirus 4 (TuAmV4) (one HHRz and one HPRz) and TuAmV1 (two HHRz). For all tested sequences, transcripts showed self-cleavage and termini of resultant fragments consistent with the predicted ribozyme cleavage-sites (fig. S3). To determine cleavage occurred *in vivo*, 5’ rapid amplification of cDNA ends (RACE) was performed from infected *Tulasnella* spp. and *Cryphonectria parasitica* extracts, and again showed perfect agreement with *in silico* and *in vitro* results (Fig. 3 A, B and fig. S4).

**Fig. 3:**
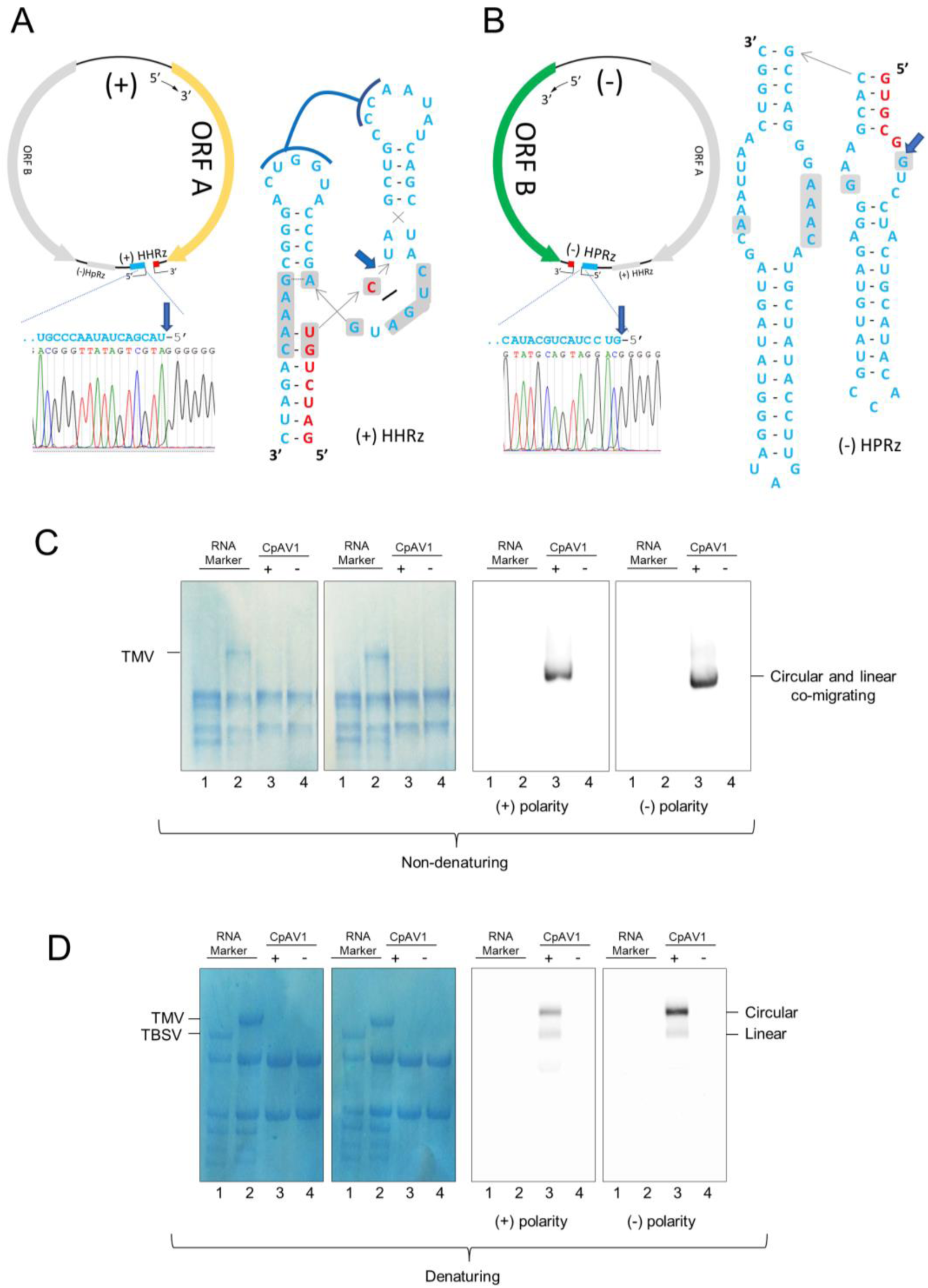
Ambiviruses encode functional +/- strand ribozymes and have circular RNA genomes. 5’ RACE and Sanger sequencing of the (**A**) positive and (**B**) negative strand of TuAmV4 genomic RNAs show *in vivo* self-cleavage (blue arrow) at the predicted cut-sites for its HHRz and HPRz motifs, respectively. Ribozyme secondary structures drawn as inlay. (**C**) and (**D**) Northern blot assays of RNA preparations separated by electrophoresis under non denaturing (C) and denaturing (D) conditions and hybridized with probes specific for the (+) and (-) CpAV1 RNA; Lanes 3 and 4 are RNAs from CpAV1-infected and non-infected *Cryphonectria parasitica*, respectively; markers are RNAs from plants, healthy or TBSV-infected (line 1 in C and D, respectively) and TMV-infected (line 2 in C and D). The single signal detected in the infected isolate with the either probe under non-denaturing conditions corresponds to both circular and linear CpAV1 genomic RNAs (4,663 nt), which are expected to co-migrate (C). In denaturing conditions (D), the ambivirus circular RNAs are delayed with respect to the respective linear forms (lower band). In C and D the two leftmost panels are total RNA stained with methylene blue. The same results were obtained from three independent experiments.

Such functional self-cleaving ribozymes on both polarities are only known to occur in diverse viroid- and Delta-like agents, all of which possess circRNA genomes which replicate through a symmetric rolling circle mechanism. Another hallmark of symmetric rolling circle replication is the *in vivo* accumulation of circRNAs of both polarities (*23*). To evaluate if ambiviruses have circular RNA genomes, northern blotting against CpAV1 was done under native and denaturing conditions (Fig. 3 C, D). Under native conditions, the CpAV1 genomic RNA migrated as a monomer (around the expected 4,623 nt), while under denaturing conditions, two bands were resolved. The retarded band would correspond to a circular molecule, and indeed the upper band shows preferential resistance to RNase R exonuclease treatment (fig. S5). Together this demonstrates that *in vivo* CpAV1, and by extension ambiviruses, replicate through a symmetric rolling circle mechanism.

Finally, to associate a phenotype to ambivirus infection, we obtained ambivirus-infected and ambivirus-free isogenic conidial isolate and show that CpAV1 causes hypovirulence in its fungal host (fig. S6), a feature that is useful for biocontrol of this important chestnut tree disease; this is the first example of a biotechnologically exploitable property linked to this new group of infectious agents. Furthermore, these agents could develop into circular RNA expression vectors, a further step into protein expression stabilization from RNA templates.

Motivated to see if other “RNA viruses” share a viroid-like genomic backbone, we searched the RDVA set for contigs with dual polarity paired-ribozymes and identified up to 16 circular contigs of approximately 3 kb encoding RdRP-like ORFs (table S2 and Fig. 4). These sequences show homology to grapevine-associated mitoviruses genomes in GenBank. Among them, we detected 7 novel circular genomes similar to *Fusarium asiaticum* mitoviruses which all carry dual-polar variants of a rare and complex self-cleaving motif, the Varkud Satellite ribozyme (VSRz), so far only described in the mitochondrial VS plasmid of some *Neurospora* isolates (*24*). The self-cleaving activity of mitovirus VSRzs was experimentally confirmed, whereas the predicted secondary structure of these agents was found not to be rod-like but highly branched (Fig. 4). These findings extend the presence of circular genomes with paired ribozymes to a different group of RNA mycoviruses and suggests that the hybridization of viroid-like agents and RNA viruses has occurred multiple times in evolutionary history.

**Fig. 4:**
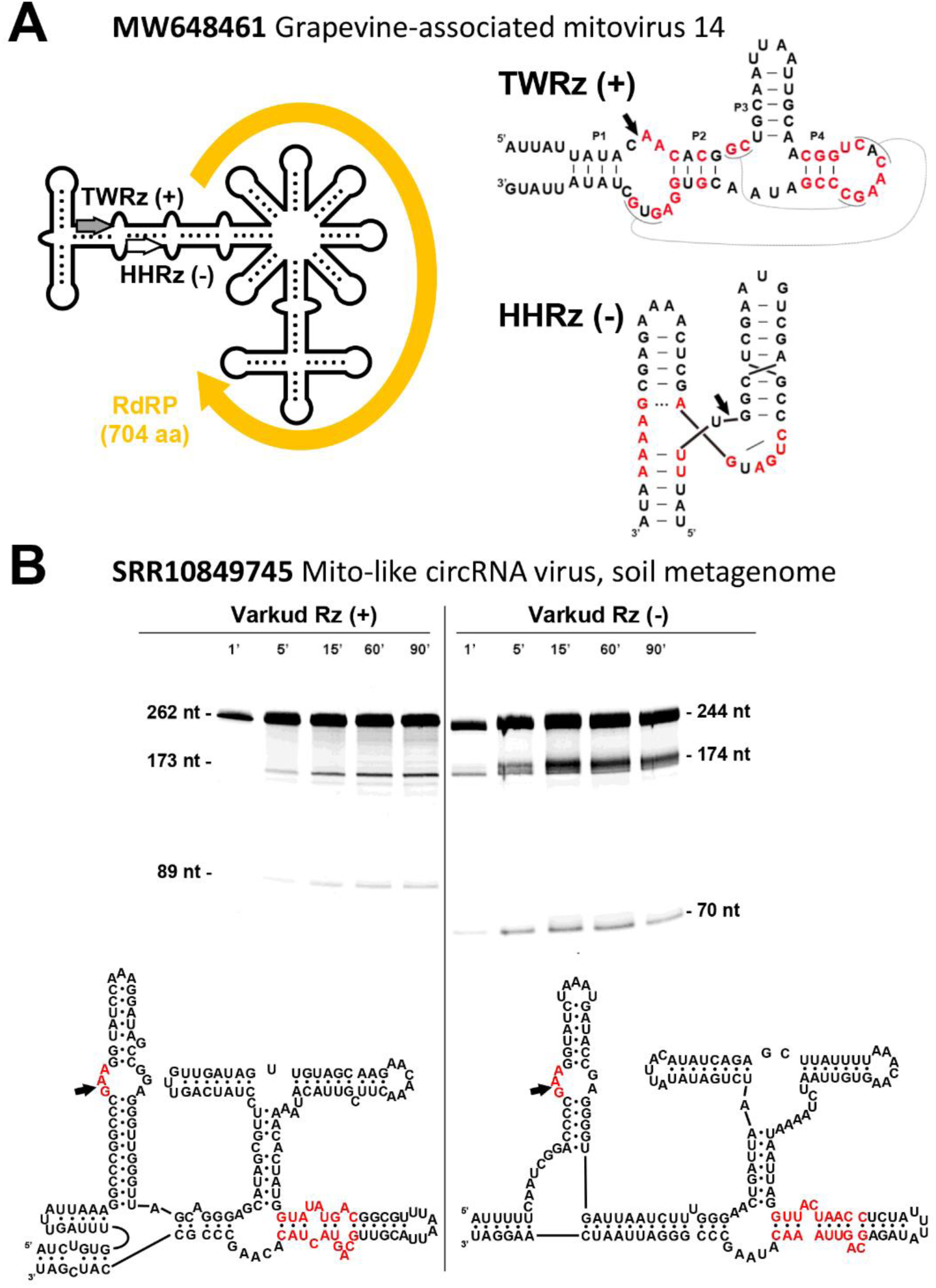
A further group of RdRP/ribozyme hybrid molecules. (**A**) Schematic genome organization of the Grapevine-associated mitovirus 14 (Accession MW648461.1). The deposited sequence contains a 203 nt repeat at both ends of the genome, indicative of a circular RNA. The predicted ORF corresponding to the RdRP is flanked by a twister (TWR) and a hammerhead (HHR) ribozyme in the plus and minus polarities, respectively. (**B**) A mitovirus with a putative circular genome detected in the library SRR10849745 (soil metagenome) show the presence of a Varkud Satellite ribozyme in each polarity. Denaturing PAGE of the RNA products from run-off transcriptions of both ribozyme constructs obtained at different incubation times shows the expected sizes of the primary and cleavage RNA products.

Infectious circular RNAs are evidently more widespread than previously appreciated, ranging from small viroid/viroid-like, medium sized delta-like, to large virus-like elements, which are hybrid forms of infectious RNAs with circular genomes encoding autocatalytic ribozymes and viral RdRPs. While computational advances will drive the exponential discovery of these elements (similar to RNA viruses) (*15, 25*), emphasis should be placed on collaborative molecular characterization of this new universe of infectious agents. While commonly we associate these entities as infectious agents of mitochondria (some mitoviruses), fungi (ambiviruses), plants (viroids), or animals (delta-like viruses), these host-distributions likely reflect our ignorance of the full diversity of infectious circular RNAs across all kingdoms.

Given the rapid mutation rate of viroids (*26*), the disposition of viroid-like elements to swap ribozymes (as shown here), and combined with the limitations of phylogenetics, the deep evolutionary relationships between infectious circRNA (such as the genomic origin of ambiviruses) is likely unknowable. Further, structurally simple ribozymes, such as HHRz can arise spontaneously in evolution (*27*), and thus embedding in relatively simple head-to-head ligated RNAs supports the notion that the simplest classes of infectious circRNA may be evolving *de novo* continuously. As viroid-like circRNA can acquire genes, such as a delta-antigen, or viral RdRP and give rise to more complex entities, it follows that host RNA polymerases may have been exapted to give rise to Earth’s primordial “viruses”. Perhaps most tantalizingly, the acquisition of an early RNA replicase ribozyme would serve as a blueprint for simplest replicator of the RNA world, and thus the origin of life.

## Acknowledgments

MdlP is grateful to Dr. Javier Forment Millet (IBMCP, UPV-CSIC) for his excellent bioinformatic assistance. This work is dedicated to the memory of Prof. Ricardo Flores who largely contributed to the study of viroids and viroid-like RNAs.

## Funding

Canadian Institutes of Health Research (CIHR) Banting Postdoctoral Fellowship #453974 (AB)

Computing resources were provided by the University of British Columbia Community Health and Wellbeing Cloud Innovation Centre, powered by AWS (AB)

Ministerio de Economía y Competitividad of Spain and FEDER grant PID2020-116008GB-I00 (MdlP)

MFA was supported by of a mobility stay grant funded by the Erasmus+ - KA1 Erasmus Mundus Joint Master’s Degree Programme of the European Commission under the PlantHealth Project.

MF is supported by a post-doctoral fellowship from the National Research Council of Italy (Assegno di Ricerca IPSP 078 2021 TO)

## Author contributions

Conceptualization: AB INZ MdlP RCE MT FDS BN

Methodology: AB MdlP RCE MT FDS MF SD MFA BN AG

Software: RCE

Formal Analysis: AB MdlP RC RCE

Investigation: AB AC MdlP RCE MF SD MFA BN AG FDS MT

Resources: SP DnA

Visualization: MdlP MF BN SD

Funding acquisition: AB MdlP MT FDS

Supervision: AB MdlP EG FDS MT

Writing – original draft: AB MdlP MT MF FDS SD BN

## Competing Interests

Authors declare that they have no competing interests.

## Data, code, and materials availability

All Serratus data are released into the public domain immediately in accordance with the International Nucleotide Sequence Database Collaboration (16) and freely available at https://github.com/ababaian/serratus/wiki/ambivirus_extended_data. Assembled genomes for this study are additionally available on GenBank under BioProject XXXXXX (pending).

Software created for this study are available at https://github.com/rcedgar/ribozy under the open-source GPLv3 license.

Biological materials, where available, are available upon request provided a MTAs is signed. Tulasnella isolate MUT4048 and MUT4047 are Deposited in Mycotheca Universitatis Taurinensis (MUT), ACP 34 is a *Cryphonectria parasitica* isolate that is being deposited in a collection and is currently available upon request to co-author DnA.

## Supplementary Materials

Materials and Methods

Supplementary Text

Figs. S1 to S6

Tables S1 to S33

## Notes

https://github.com/rcedgar/ribozy

https://github.com/ababaian/serratus/wiki/ambivirus_extended_data

### Competing Interest Statement

The authors have declared no competing interest.

