## Supplementary Materials for "Extant hybrids of RNA viruses and viroid-like elements"


**This PDF file includes:**

Materials and Methods

Supplementary Text

Fig. S1 to S6

Table S1 to S3

### Materials and methods

Ribozyme search of the Sequence Read Archive

Observing that ribozymes are sufficiently short to be captured on a short sequence read (less than 100 nt), we reasoned it will be possible to screen large volumes of sequencing data to identify libraries potentially containing ribozyme agents. To this end we adapted the ultra-high throughput Serratus cloud computing platform (*15*), with the INFERNAL (*14*), an algorithm for identifying RNA elements using secondary structure and nucleotide covariation modelling. Using Serratus-INFERNAL (https://github.com/ababaian/serratus/tree/infernal-dev), we performed a pilot screen of 198,194 publicly available metagenomic/metatranscriptomic sequencing libraries in the Sequence Read Archive (SRA) for the delta, hairpin, hammerhead, twister, and Varkud satellite ribozymes. The SRA search query was randomly sub-sampled from the set of “"METAGENOME" OR "METATRANSCRIPTOME" OR "metatranscriptomic"[Filter] OR "metagenomic"[Filter] NOT amplicon [All Fields] AND "platform illumina"[Properties] AND cluster_public[prop]” on 2022-01-13. We then heuristically rank ordered the libraries based on the quality and number of distinct ribozyme hits in a library as well as sequencing depth for focused sequence assembly of the SRA. Candidate viroid-like and ribozyviria sequences were identified by the following criteria, all of which must be satisifed: (1) the presence of 5’-3’ end k-mer overlaps suggestive of a circular molecule, (2) a pair of complementary hits to one ribozyme model with E-value < 1e-3, and (3) < 25% repetitive sequence according to the union of hits reported by dust (*28*), Tandem Repeat Finder (*29*) and self-hits according to ublast (*30*). Clustering into 90% identity species-like Operational Taxonomic Units was performed using circuclust (https://github.com/rcedgar/circuclust).

Bioinformatic detection and phylogeny of ambiviral RdRPs

RdRPs (ORF-A) from either the 19 deposited Ambivirus genomes at the GenBank (09-2021), or the 169 ambiviral-like sequences (either putative full circles or genomic fragments) detected though INFERNAL in unrestricted/public metatranscriptomic datasets of the JGI database ("metatranscriptome" OR "metatranscriptomic"[Filter]), were used to build a consensus sequence for the ambivirus motifs A, B and C. Based on the obtained consensus sequence for the ambiviral palm domain, we build a new version of the palmscan software (https://github.com/rcedgar/palmscan) capable of ambivirus RdRP detection. Palmscan searches were performed in the circular contigs from the RDVA, which together with the ones obtained from GenBank and JGI databases resulted in 439 ambiviral RdRPs at 90% sequence identity (https://github.com/ababaian/serratus/wiki/ambivirus_extended_data/). We performed multiple sequence alignment on the clustered RdRP palmprint sequences using MUSCLE (*31*) and maximum-likelihood tree is inferred from the alignment using MEGA11 (*32*). Secondary RNA structures of minimum free energy of the ambivirus genomes were calculated with the RNAfold program from the ViennaRNA Package (*33*). The obtained structures were visualized by circular plots obtained with the jupiter software (https://github.com/rcedgar/jupiter)

Fungal growth in liquid media

To obtain fungal mycelia for molecular analysis, three CpAV1-free and three CpAV1-infected isolates were inoculated in liquid media in biological triplicate (18 samples). The media contained a mix of 24 g/L potato dextrose broth (PDB) (SIGMA) with two European chestnut (*Castanea sativa*) sticks of 8cm length (cut longitudinally) autoclaved at 121 $^{o}$C for 1 hour. 50 mL of the mix was transferred into 100 mL conical flasks for all the samples. Then, 12.5 mL of the liquid media was collected in 50 mL conical Falcon$^{TM}$ tubes (Corning$^{TM}$ Fisher Scientific) where 30 mycelia plugs were transferred for each of the replicates. The tissues were ruptured with Omni Tissue Homogenizer (Qiagen) for 1 minute and 3.5 mL of the homogenized paste was inoculated in each of the conical flasks containing the liquid media. The cultures were kept constantly shaking at 120 rpm at room temperature.

At three days post-inoculation, the fungal cultures were harvested by filtering under vacuum in a Buchner funnel covered with one layer of Miracloth (Calbiochem) in a vacuum pump to separate the mycelia from the liquid media, the harvested mycelia were frozen at -80 °C and lyophilized for 45 hours (EDWARDS MODULYO freeze dryer). Mycelial dry weight for each replicate was determined with a top pan balance.

*Tulasnella* spp isolates were grown in liquid cultures and harvested and lyophilized as above.

RNA extraction

RNA extraction was done with Spectrum Plant Total RNA kit, (SIGMA-Aldrich) following manufacturer's protocol with minor modifications: briefly, lyophilized mycelia (100 mg) was broken in 1.5 mL conical microcentrifuge tubes with O-rings (Biosigma) with 10, 4.5 mm diameter, ceramic beads in a FastPrep-24$^{TM}$ homogenizer from MP Biomedicals$^{TM}$ for 30 seconds at setting 6. A 1 mL mixture of lysis buffer and 2-mercaptoethanol (Sigma-Aldrich) at 1: 100 volume dilution was added to the samples and mycelia was extracted by another round of bead beater, as above; the sample was then centrifuged at 10,000 g for 4 minutes. 700 µL of the supernatant was then transferred to filter column tubes provided in the kit and centrifuged for 30 seconds at the same speed. The columns were discarded and 700 µL of the binding solution was added to the flow-through. This mixture was then passed through the binding column by a further centrifugation as above. The column was collected, transferred to a fresh collection tube and washed twice with wash buffer I and II (600 and 700 µL respectively). RNA was then eluted in a 40 µL elution buffer. RNA quantification was done with Nanodrop LITE Spectrophotometer (Thermo Fisher Scientific).

Characterization of terminal sequences of ambivirus genomes *in vivo*

Rapid Amplification of cDNA Ends (RACE) analysis was performed on CpAV1 from *C. parasitica* strain ACP34 (*13*) and TuAmV1 and TuAmV4 from Tulasnella spp. (respectively from strains MUT4048 and MUT4047) (*12*) using the Hirzmann method (*34*). Briefly, cDNA of the 5 ʾend of the positive and negative sense of the ambivirus genome was synthesized with the First Strand cDNA synthesis Kit #K1612 from Thermo Scientific with a specific primer (table S3) following manufacturer's protocol, and cleaned with Zymo clean and concentrator 5 kit (Zymo research, Irvine, CA, USA) and treated with Terminal Transferase (NEB, Ipswich, MA, USA) to add a poly-A or poly-G tail. After terminal transferase reaction, the cDNAs were cleaned using DNA Clean & Concentrator™ 25 (Zymo research, Irvine, CA, USA) with a final elution volume of 10 µL and 1 µL of the obtained cDNAs was used as template to perform a PCR reaction with Taq polymerase (OneTaq kit from Biorad) following manufacturer's protocol using a specific primer and anchored poly-T or poly-C primer complementary to the added tail at 51° C in the annealing step for 30 sec, 15 sec at 68° C for extension step and 98°C for the denaturation step for 15 seconds. The resulting amplicons were fractionated by gel electrophoresis in 1% Agarose TAE buffer and purified using a Zymo gel DNA recovery kit (Zymo research, Irvine, CA, USA) and cloned in pGEM-T easy vector system (Promega, Madison, WI, United States) following manufacturer's instructions. Ligation products were transformed in DH5 alpha E. coli cells prepared and transformed following exactly the protocols and materials of the Mix&Go! E.coli Transformation Kit (Zymo Research). Sanger sequencing of the obtained clones (at least 3 clones containing the terminal region, for each construct) was performed by Biofab research (Rome, Italy) using the specific primers detailed in table S3.

Attempts were performed to directly sequence the 3 ʾend of both the positive and negative sense of the CpAV1 and TuAmV1 ambivirus genome by a method modified from Lambden et al. (doi: 0022-538X/92/031817-06$02.00/0), which is optimized for the sequencing of double stranded RNA, but all failed. Briefly, a 5ʾ-phosphorylated, 3ʾ-amino-linked oligodeoxy-nucleotide primer was ligated to the 3 ʾtermini of the total extracted RNA (1 µg total RNA) with T4 RNA ligase at 4° C over-night. The ligated RNA was cleaned in column (binding column of the Spectrum Plant Total RNA kit, SIGMA), the cDNA was synthetized (Superscript IV, Thermo-Fisher) with a primer complementary to the ligated adaptor, cleaned using Zymo clean and concentration 25 kit (Zymo research, Irvine, CA, USA) with a final elution volume of 25 µL; 1 µL of the obtained cDNAs was used to perform a PCR reaction using a specific primer and the primer used for the cDNA synthesis with the same conditions described above, except the annealing temperature at 55° C.

Analyses of RNA self-cleavage *in vitro*

Self-cleavage activity of HHRz and HPRZ of representative ambiviruses (CpAV1, TuAmV1 and TuAmV4) and VSRz of the mito-like virus detected in the SRR10849745 library was tested *in* *vitro* by transcription of recombinant plasmids containing the cDNA of the respective ribozymes. The cDNAs of viral RNA fragments containing the ribozymes were either amplified using the appropriate primers listed in table S3 or purchased as gBlock Gene Fragments (Integrated DNA Technologies), subcloned in either pBlueScript KS or pGEM-T easy Promega, (Madison, WI, United States) and sequenced to confirm the orientation and absence of unwanted mutations. The recombinant plasmids, linearized with the appropriate restriction enzyme, were transcribed *in vitro* with T7 RNA polymerase (Thermo-Fisher Scientific) at 37° C for 2hr using buffer provided by the manufacturer and the reaction products separated by denaturing 5% PAGE 8 M urea and 1xTBE (89 mM Tris, 89 mM boric acid, 2.5 mM EDTA, pH 8.3), stained with ethidium bromide and UV visualized. The 5 ʼterminal nucleotide of the 3 ʼself-cleavage product of HHR and HPRz motifs was determined by 5 ʼRACE experiments. Briefly, the 3 ʼRNA fragment generated by the self-cleavage of each tested ribozyme during the *in vitro* transcription was eluted from the gel by grinding with a mixture (1:1) of water-saturated phenol and buffer (100 mM Tris-HCl pH 8.9, 1 mM EDTA, 0.5% SDS), and recovering the nucleic acid from the aqueous phase by ethanol precipitation (*35*) then their 5 ʼterminal sequence was determined by 5ʼRACE as described above using the primers indicated in table S3).

Northern-blot hybridization assays

Agarose (1%) gel electrophoresis under denaturing conditions of total RNAs was performed in denaturing condition using 6 M glyoxal and 50% v/v DMSO in HEPES EDTA buffer, keeping the RNA samples at 55 °C for 20 minutes prior to loading, following the protocol described previously (*12*). Agarose gel electrophoresis of RNA preparations under non-denaturing conditions was carried out in 1X TAE without denaturation prior to loading.

After electrophoresis, RNA was blotted on Immobilon-Ny+ membrane (Merck, Darmstadt, Germany) and crosslink was performed in a Biorad gs gene linker UV chamber (Biorad, Hercules, CA, USA). The membranes were hybridized with digoxigenin (DIG)-labeled riboprobes specific for (+) or (-) polarity strands of CpAV1 that were synthesized by *in vitro* transcription using the Dig-RNA labeling mix (Roche Diagnostics GmbH, Germany) and linearized plasmids containing CpAV1 cDNA inserts from positions 2798 to 3088 (located in ORFA) and from positions 1628 to 1998 (located in ORF B), respectively. Pre-hybridization and hybridization were performed in DIG easy Hyb (Roche Applied Science, Germany) buffer at 65 ◦C. After hybridization, the membranes were washed twice in 2× SSC (0.3 M NaCl, 0.03 M sodium citrate, pH 7), 0.1% SDS at room temperature for 10 min, and twice in 0.1× SSC, 0.1% SDS at 65◦ C for 15 min. The hybridization signals were revealed with the anti-DIG alkaline phosphatase conjugate AB fragments and the chemiluminescence substrate CDP-star (Roche Applied Science, Germany) following the manufacturer instructions and visualized with a Chemidoc Touch Imaging system (Bio-Rad, Hercules, CA, USA).

qPCR on the terminal sequences of the cleaved genome

In order to investigate whether the terminal fragments of the genomic and antigenomic strands of the ambivirus RNAs were fully complementary, we performed strand-specific qPCR analysis on TuAmV1 RNAs. cDNAs were synthesized from the same amount of RNA with primers specifically annealing either on the genomic or on the antigenomic strand and (table S3) designed either on the terminal putative sequence found by RACE, or on a region up/downstream. A pair of primers was designed on the ORF A and the qPCR product obtained was used as internal reference for the quantification of the terminal sequences. The cDNAs were column-purified (DNA Clean & Concentrator™ 25 Zymo Research) and a standard qPCR was performed using the same primers: briefly, qPCR reaction was performed in 10 µL final volume using the SYBR® Green Master Mix (Biorad). Each 10 µL reaction mix contained 5 µL mix, 4 µL of sterile water and 0.2 µL of forward and reverse primers (Supplementary table S3), and 1 µL of cDNA. Amplifications were carried out in 96-well plates in a CFX Connect Real-Time PCR Detection System (Biorad) with thermocycling conditions of 3 min at 95°C, 20 seconds at 95°C, and 30 seconds at 60°C for 40 cycles. The quantification was reported as quantity relative to the internal reference.

RNAse R digestion

RNAse R (Lucigen) was used to test the circular nature of the ambivirus genomes on CpAV1. Samples containing 2 µg of total RNA from infected *C. parasitica* ACP43 were incubated in RNAse R 1X reaction buffer, 1 U of RNAse R for 5, 15 or 30 minutes at 37°C. Negative controls were obtained mixing 2 µg of total RNA with the reaction buffer without RNaseR and tested at time 0 and after 30 minutes of incubation at 37°C. Reaction inactivation was performed adding the northern blot denaturation mix (see northern description above) to each samples and incubating at 65°C for 20 minutes. Denatured samples were then analyzed through denaturing agarose gel electrophoresis and northern blotting as reported above.

Deriving ambivirus-infected and ambivirus-free isogenic lines

Isolate -ACP34- originally collected from a diseased tree (*Castanea sativa*) with evident cankers in Azerbaijan (*13*). The *C. parasitica* isolates were maintained on PDA media at 4 °C.

Isolate ACP34 was inoculated in 90 mm Petri plates with PDA for two months and sixteen days with light/dark cycles of 12 hours. Conidia were harvested from the culture by pipetting 10ml of sterile water in the Petri plate for two minutes and scrubbing the mycelia twice at one-minute intervals. The spore suspension aliquot was collected and filtered with cotton placed in a 5 mL tip. Isogenic monoconidial isolates were obtained by serially diluting the harvested spore suspension and plating 300 mL of the diluted conidia in 150 mm Petri plates with serial dilutions (1:$10^{-1}$ to 1:$10^{-10}$). Single spore isolates were transferred after 5 days, and single fungal colonies were tested for ambivirus presence/absence by qRT-PCR. cDNA was synthesized with the High Capacity cDNA Reverse Transcription Kit (Appliedbiosystems, by Thermo Fisher Scientific) following manufacturer's protocol.

To diagnose CpAV1 in the isolates, qRT-PCR was performed using a 1700 fast SDS Real-Time PCR detection system (Applied Biosystems). The PCR reaction was performed in 10 µL using the I-Taq supermix (Biorad) and Taqman probe. Each 10 µL reaction mix contained 5 µL supermix, 0.15 µL probe, 4 µL of sterile water and 0.2 µL of forward and reverse primers (Supplementary Table S3), and 1 µL of cDNA. Amplifications were carried out in 96-well plates in a CFX Connect Real-Time PCR Detection System (Biorad) with thermocycling conditions of 3 min at 95$^{o}$C, 20 seconds at 95$^{o}$C, and 30 seconds at 60$^{o}$C for 40 cycles.

*In vitro* Culture on Potato Dextrose Agar

After deriving the isogenic CpAV1-free and CpAV1-infected isolates, mycelia plugs were taken from the edges of subcultures of three isolates for each condition (+1, +4, +8, -2, -7, -12). The isolates were inoculated in 150 mm Petri plates and incubated in potato dextrose agar (PDA) (Sigma-Aldrich) at 26°C for 14 days. For each isolate we included three biological replicates and mycelia diameters were measured for all isolates at three days intervals post-inoculation until the full diameter of the Petri plate is covered.

Virulence characterization of virus-infected and virus-free isogenic isolates on live chestnut stem and apple fruits

Live chestnut stems of 30 cm were prepared for inoculating the isogenic CpAV1-free and infected isolates (1+, 4+, 8+, 2-, 7-, 12-). Cuttings to be used in the experiment were obtained from chestnut suckers from a chestnut orchard (European chestnut, *Castanea sativa* in Salò-BS, Italy). Each stem cutting was disinfected with 70% ethanol then the cuttings were coinfected with virus-positive and negative mycelia plugs at a 17 cm distance from each other. The experiment was performed with three biological replicates randomly distributed to have CpAV1-infected and CpAV1-free mycelia plug at the upper and lower side of the stems. A cork borer of 5 mm diameter was used to artificially displace a section of the bark. Mycelia plugs were then placed under the removed barks and tightly taped for wound closure. Also, negative controls without mycelia plugs but with similar wounds were made. The cuttings were stored in 1.5 L of vermiculite soaked with water for 21 days.

Apples (cv Delicius) were also used to verify virulence. Each apple was inoculated in 4 points with a CpAV1 infected and a CpAV1 free strain (2 replicates each on a single apple, in opposite positions). Inoculated apples were kept at room temperature for 14 days.

Statistical Analysis

Analyses of the data referred to the measures of lesion or canker size in the virulence assays (apples and chestnut cuttings) were performed with one-way analysis of variance (ANOVA) using the R-statistical program (*36*). Significant differences were assessed with Tukey's posthoc test ($p<0.05$).

### Supplementary Text

The linear form of the positive sense and negative sense genomes of ambiviruses are not complementary along the full-length segments

All the Baltimore classes of RNA viruses (III, IV and V) contain viruses that replicate via RNA dependent RNA polymerases: in these classes these enzymes copy the full-length template linear genomic RNA in a full length antigenomic linear RNA for maintaining genomic RNA identity during replication. Presence of complementary full length genomic and antigenomic RNAs is, to our knowledge, without exception in RNA viruses and is a hallmark of their replication strategy. This implies that the 5 'end of the antigenomic RNA is complementary to the 3 'end of the genomic RNA, and often, when antigenomic RNA accumulation is abundant, the nt sequence of the 3 'end of the genomic RNA can be inferred indirectly by sequencing the 5 'end of the antigenomic RNA with Hirzmann's approach (*34*) .

Given that ambiviruses accumulate in fungi as virion-less entities, genomic and antigenomic RNAs are purely conventional and we follow a previously proposed convention where the positive sense (Genomic) RNA is the one encoding for ORF-A, the putative RdRP, while antigenomic (negative) sense RNA is the one encoding for ORF-B (*12*). Given these premises, we performed two distinct 5 'RACE experiments using oligonucleotides close to the presumed 5 ' end of the genomic and antigenomic linear monomeric RNAs on three distinct ambiviruses (TuAmV1, TuAmV4 and CpAV1) in total RNA from infected fungal extracts (*Tulasnella spp.* and *Cryphonectria parasitica*) and in all three cases we had surprising results: the first 245 nt for CpAV1, 256nt for TuAmV4 and 223nt for TuAmV1 of the 5 'of the genomic RNAs were exactly complementary to : the first 245 nt for CpAV1, 256nt for TuAmV4 and 223nt for TuAmV1 5 'of the antigenomic RNA. If genomic and antigenomic would be complementary along the full length, this would imply that the monomeric genomic and antigenomic have a same orientation sequence repeat of circa 250 nt (fig. S1 A), which would be unprecedented for RNA viruses. This prompted us to further investigate if such repeat indeed exists on the genomic and antigenomic RNA-

Attempts to sequence the 3 'ends of ambivirus RNAs of both polarity strands failed, which is consistent with the cleavage by HHRz and HPRz motifs which leave a blocking 2'-3 'cylic phosphate on the 3 'termini (*37*).

Given that we failed to use alternative RACE methodology to determine more directly the 3 ' end of the genomic RNA, we proceeded to carry out RT-PCR amplification, that would use oligonucleotides to prime cDNA synthesis specific for the genomic or antigenomic RNAs. We selected for RT-qPCR amplification regions of the genome across the putative link between the conserved repeat and the specific sequences upstream (for determining the presence of the 3 ' putative repeat in the genomic RNA). As control, we included in the same assay the quantification of the central region of the RNA genomic segment. Surprisingly, the ratio between the two is not consistent with the expected equimolar existence of the two repeats at the 5 'and 3 'end of the genomic RNA (fig. S1 B).

The same experiment carried out for the antigenomic RNA provided the same results: the repeat at the 3 'end is not present in the expected same concentration as the 5 'end (fig S2, panel C). Minimal amount of the repeats are indeed present because in the same extracts we have shown accumulation of putative dimer or circular RNAs of both strands, that would indeed provide some amplification.

In conclusion, the two monomeric forms of the genomic and antigenomic RNA are not full-length complementary RNA, pointing to a replication mechanism for these RdRP encoding RNAs different from that of the other Baltimore classes of RNA viruses.

These data are consistent with the different positions of the ribozyme self-cleaving sites in the genomic and antigenomic RNAs.

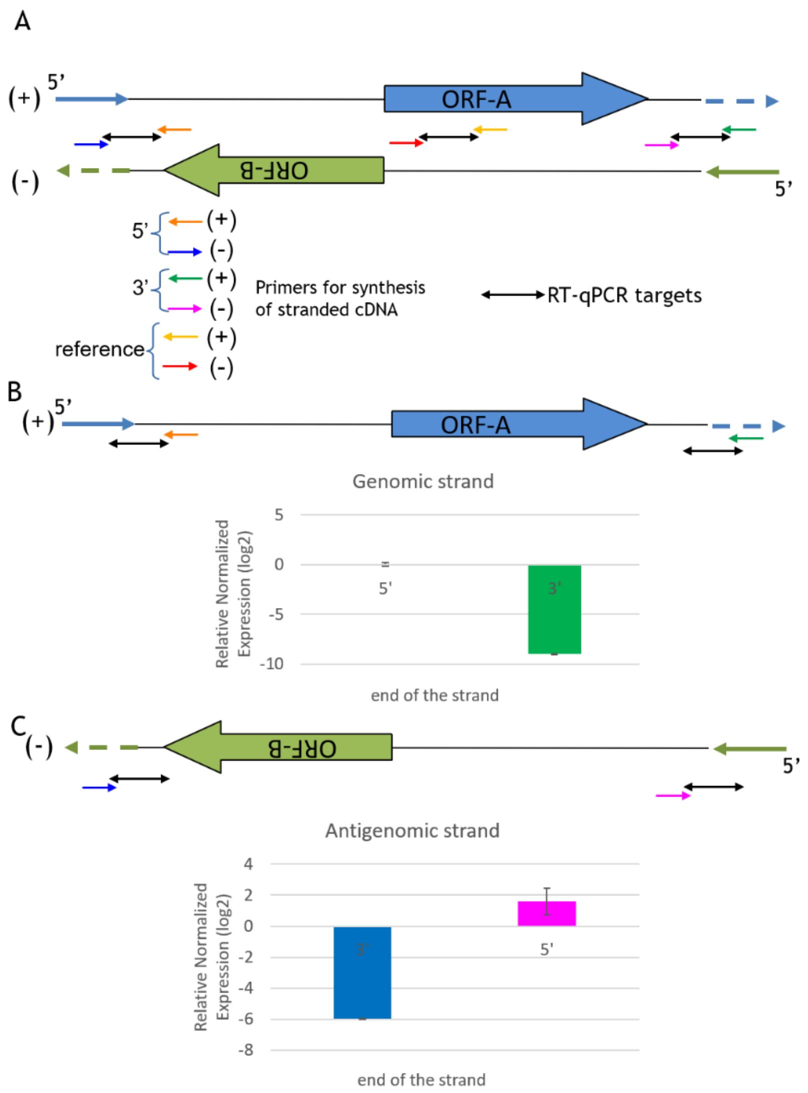


**Fig. S1. The linear replication intermediates of both polarity strands are asymmetrical**

(**A**): RACE analysis of 5 'ends of genomic (+) and antigenomic (-) RNA of TuAmV1 revealed 222 nt complementary sequences (thin solid arrows): broken arrows correspond to putative same sense repeats in the genomic and antigenomic segment. Thinnest colored and black arrows respectively correspond to strand specific oligonucleotides and to amplification products across the putative junctions between the sequence repeats and the specific region of the genomic and antigenomic RNA, or on the ORF-A coding sequence (red and yellow arrows) as an internal reference for the abundance of the genomic and antigenomic strands. (**B**): relative quantification (log2) of the genomic strand RNA region across the junction of the 5 'and 3 'repeat. (**C**): relative quantification (log2) of the antigenomic strand RNA region across the junction of the 5 'and 3 'repeat. The color of the bars reflects the oligonucleotide used for the cDNA synthesis and represented by the thinnest arrows. In the 5 'end region of the genomic, the accumulation orders of magnitude larger than the putative 3 'end of the genomic RNA. For the antigenomic RNA the 5 'end accumulates more than the 3 'end. For both the genomic and the antigenomic strands the accumulation of the 5 'region is comparable to the one of the internal reference region on the ORF-A.

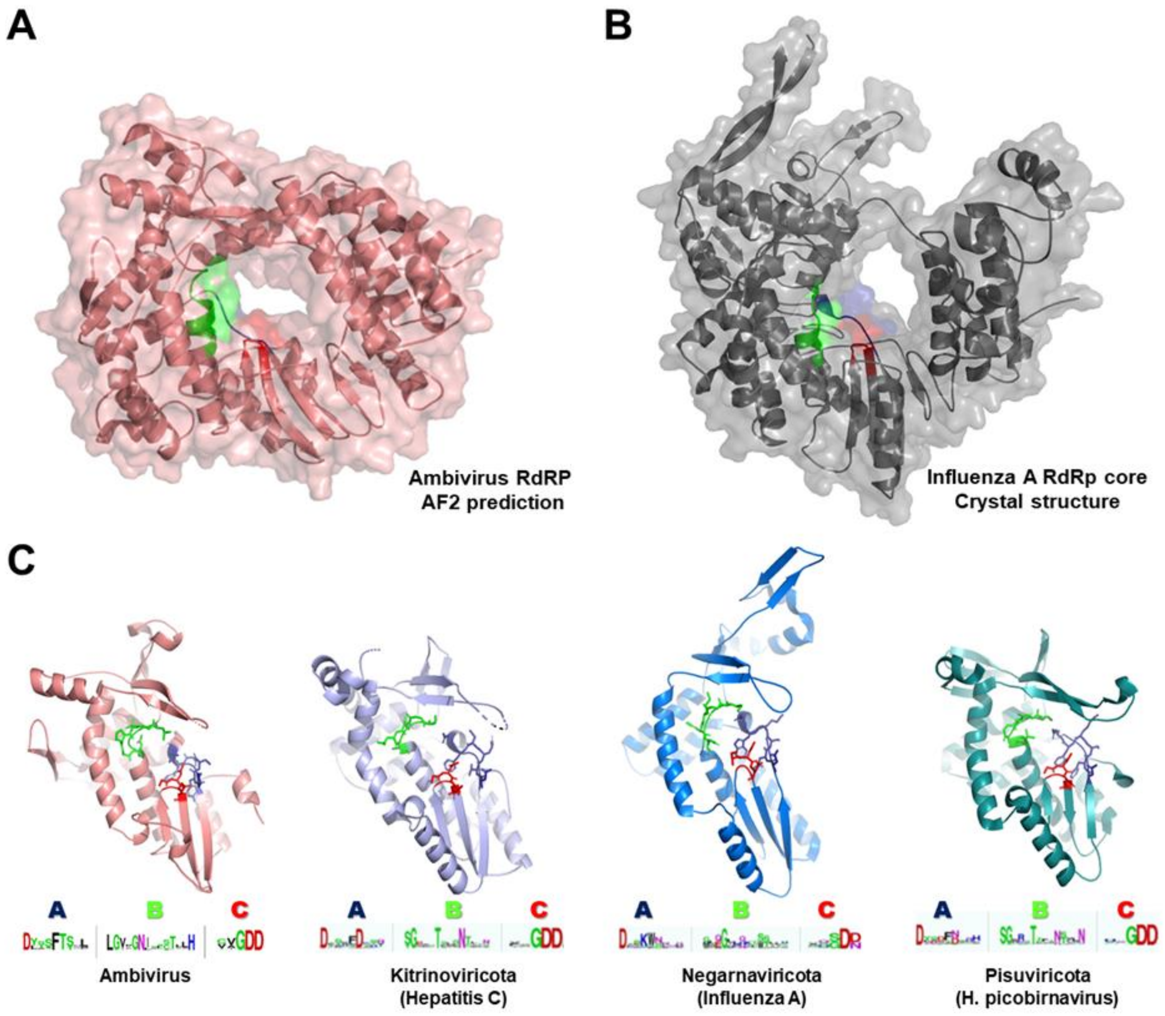


**Fig. S2. Structural models for the RNA-dependent RNA polymerases**

Structural models for the RNA-dependent RNA polymerases (RdRP) of (**A**) the *Armillaria borealis* ambivirus 2 (Accession MW4238010.1, as predicted by AlphaFold2 algorithm) and (**B)** the bat influenza A RdRP core (PDB 6t0u, chain B, as obtained by cryoelectron microscopy (38) show a high global similarity (Z score 19.2, rmsd 4.0 A) and equivalent placement of the putative catalytic residues of the palm motif (A box in blue, B box in green and C box in red). (**C)**. Structural comparison of the RdRP palm domains of the *Armillaria borealis* ambivirus 2 (Alphafold2 predicted, see A) and those experimentally determined for Hepatitis C (PDB 3br9), Influenza A (PDB 4wsb) and human picobirnavirus (PDB 5i61) RdRPs. Highly conserved catalytic residues for each palm motif are shown as sticks. Consensus sequences of the motifs A, B and C for each viral phylum are shown at the bottom.

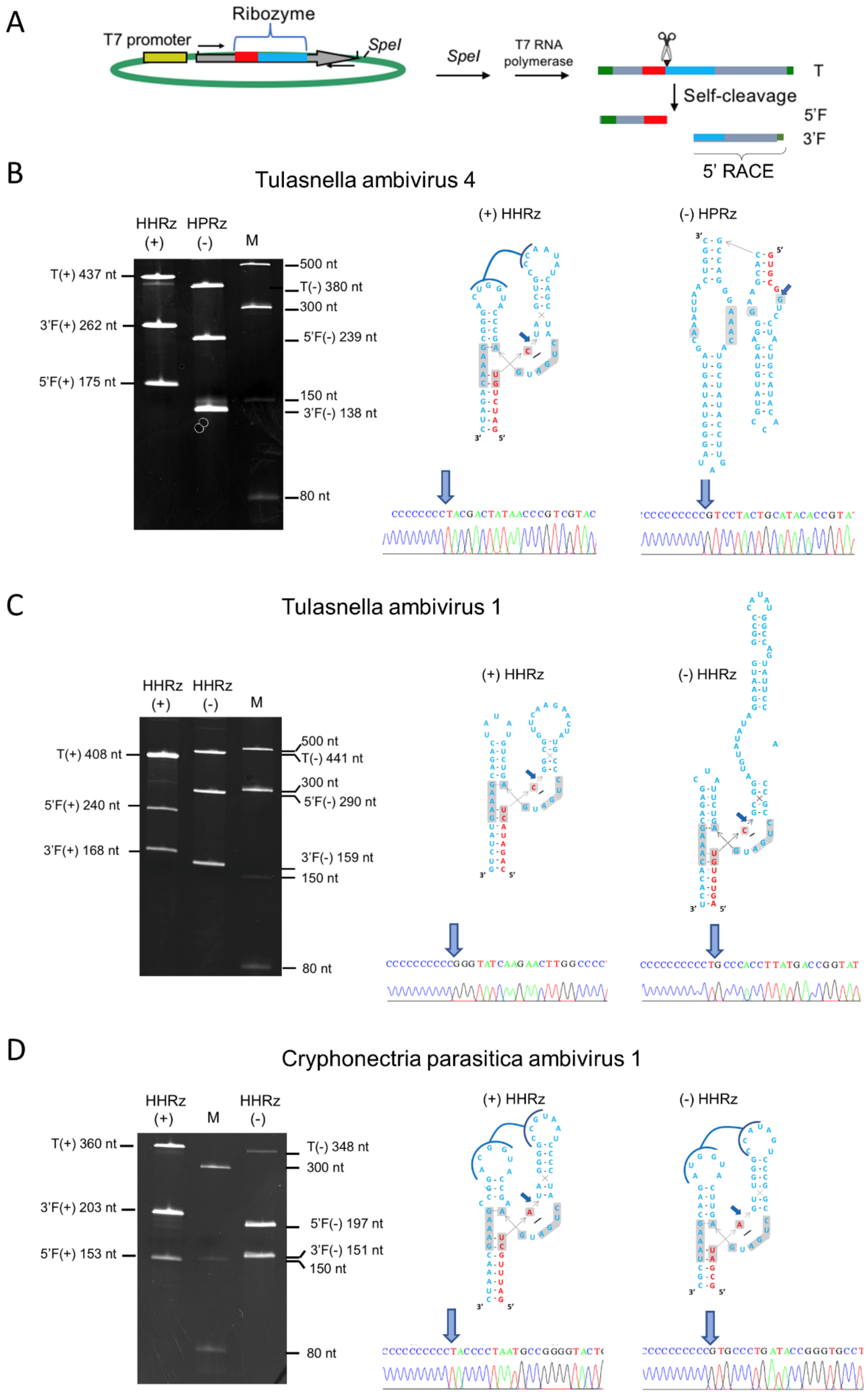

A
Ribozyme
T7 promoter
SpeI
SpeI
T7 RNA polymerase
T
Self-cleavage
5'F
3'F
5' RACE
B
Tulasnella ambivirus 4
HHRz (+)
HPRz (-)
M
T(+) 437 nt
3'F(+) 262 nt
5'F(+) 175 nt
500 nt
T(-) 380 nt
300 nt
5'F(-) 239 nt
150 nt
3'F(-) 138 nt
80 nt
(+) HHRz
(-) HPRz
C
Tulasnella ambivirus 1
HHRz (+)
HHRz (-)
M
T(+) 408 nt
5'F(+) 240 nt
3'F(+) 168 nt
500 nt
T(-) 441 nt
300 nt
5'F(-) 290 nt
3'F(-) 159 nt
150 nt
80 nt
(+) HHRz
(-) HHRz
D
Cryphonectria parasitica ambivirus 1
HHRz (+)
M
HHRz (-)
T(+) 360 nt
3'F(+) 203 nt
5'F(+) 153 nt
T(-) 348 nt
300 nt
5'F(-) 197 nt
3'F(-) 151 nt
150 nt
80 nt
(+) HHRz
(-) HHRz

**Fig. S3. Ribozymes contained in ambivirus RNAs are active *in vitro.* (A)** Schematic representation of the DNA template for the *in vitro* transcription, consisting of a plasmid containing the cDNA of the self-cleaving ribozyme to be tested, and of the expected RNA products. Plasmids containing the ribozyme sequences of Tulasnella ambivirus 4 (TuAmV4), Tulasnella ambivirus 1 (TuAmV1) and Cryphonectria parasitica ambivirus (CpAV1) were linearized and transcribed with T7 RNA polymerase to produce full-length transcripts (T) and the respective 5' and 3' fragments (5'F and 3'F, respectively) derived from the ribozyme self-cleaving activity. In green, plasmid sequences; in yellow, polymerase promoter; in grey ambivirus RNA sequences with 3'F and 5'F ribozymatic regions denoted in blue and red, respectively. The predicted self-cleavage site is indicated by a arrowhead. **(B-D)** Left panels, analyses by denaturing 5% PAGE of the *in vitro* transcription products of the plasmids containing the cDNAs of the self-cleaving ribozymes identified in the (+) and (-) polarity strands of the genomic RNAs of TuAmV4, TuAmV1 and CpAV1. The sizes of the full-length, 3'F and 5'F RNAs generated during each transcription were consistent the self-cleaving activity of the ribozyme in each transcript. M, RNA ladder with sizes indicated on the left; numbers on the right and on the left indicate the size of the RNAs. Middle and right panels, primary and secondary structure of (+) and (-) ribozymes contained in the viral genomic RNAs, respectively, of TuAmV4 (B), TuAmV1 (C) and CpAV1 (D). Self-cleavage site of each ribozyme was confirmed by 5' RACE of the 3'F fragment, with the respective sequencing electropherograms reported on the bottom. The predicted self-cleavage site is indicated by a blue arrow. The nucleotides of the catalytic core conserved in most natural HHRz and HPRz structures are reported on a grey background.

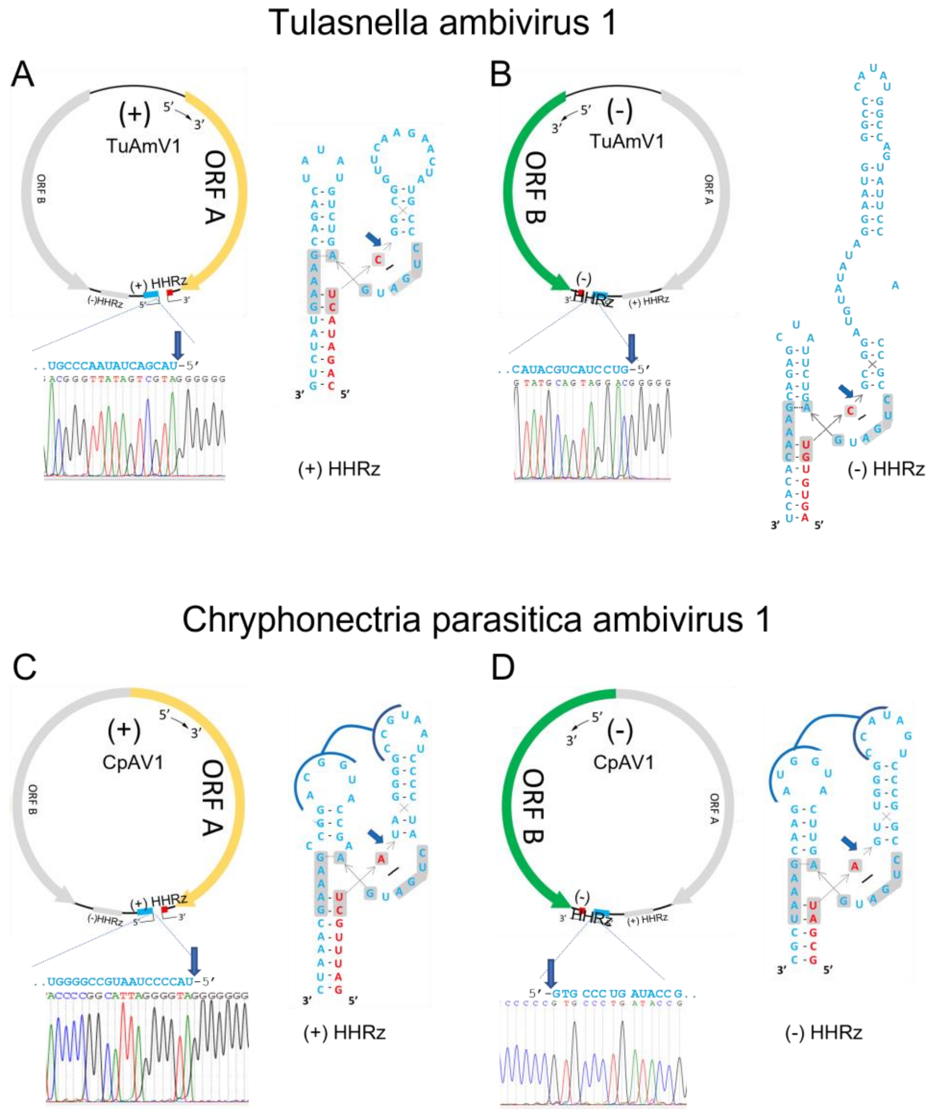


**Fig. S4. The 5′ terminal end of (+) and (-) genomic RNAs of both polarity strands of Tulasnella ambivirus 1 (TuAmV1) and Cryphonectria parasitica ambivirus 1 (CpAV1) are coincident with those predicted to be generated by the RNA self-cleavage mediated by the respective encoded ribozymes.** (**A** and **B**) refer to (+) and (-) polarity strands of the genomic RNA of TuAmV1, respectively. (**C** and **D**) refer to (+) and (-) polarity strands of the genomic RNA of CpAV1, respectively. In each Panel: Left, up, schematic representation of the (+) (A and C) and the (-) (B and D) genomic RNAs of TuAmV1 and CpAV1, respectively, with the encoded ORF A and HHRz depicted in yellow and red/light-blue for the (+), and the encoded ORF B and HHRz depicted in green and red/light-blue for the (-) polarity strand; the position of the ORF and the ribozyme contained in the opposite polarity are reported in grey. Left bottom, electropherogram of the sequenced 5′ RACE products showing the 5′ terminal end of each viral RNA, which is coincident with the one predicted based on the self-cleavage mediated by the respective encoded HHRz. Right: secondary structure of the HHRz contained in each viral RNA polarity strand; the predicted self-cleavage site is indicated by a blue arrow with the sequences at the resulting 3′ and 5′ terminal ends depicted in red and light blue, respectively. Nucleotides conserved in most natural HHRzs are reported on a grey background.

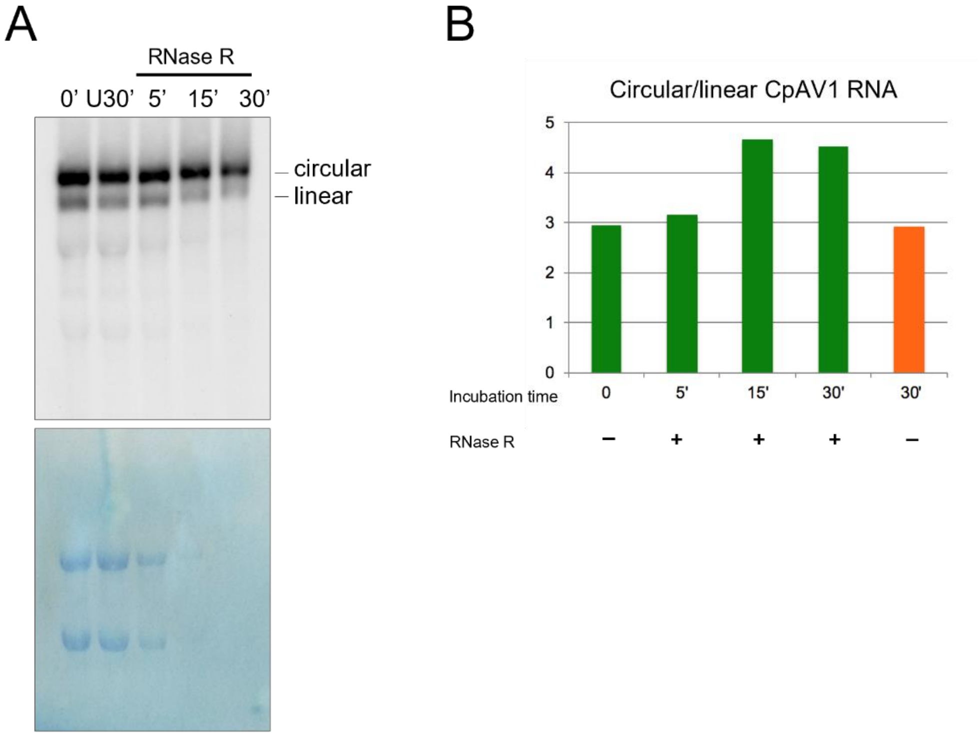


**Fig. S5. Circularity of ambivirus RNAs assessed by RNase R treatment.** RNA preparations (2 ug) from *Cryphonectria parasitica* infected by CpAV1 were incubated with RNase R (1 UNIT) during 5, 15 and 30 min at 37°C in buffer supplied by the manufacturer. Controls at time 0 and at 30 min were incubated in the same conditions without RNase R. (**A**) Northern blot hybridization of treated RNA preparations with a specific Dig-RNA probe detecting the (-) polarity of CpAV1 (upper panel). Ribosomal RNAs were visualized on the membrane by toluidine blue staining (bottom Panel). While linear forms were almost completely degraded after incubation with RNase R for 30 min, the circular forms clearly resisted. No major changes of both the linear and circular RNA forms were observed after 30 min of incubation in the absence of RNase R. (**B**) Relative accumulation of circular/linear RNA forms of CpAV1 at different incubation times with RNase R based on the quantification of the Northern-blot hybridization signals. While in the absence of RNase R the ratio circular/linear RNAs does not change after 30 min incubation time (orange bar), such a ratio increases in the presence of RNase R at the different time points, showing a higher resistance of the circular forms. The same results were obtained from three independent experiments.

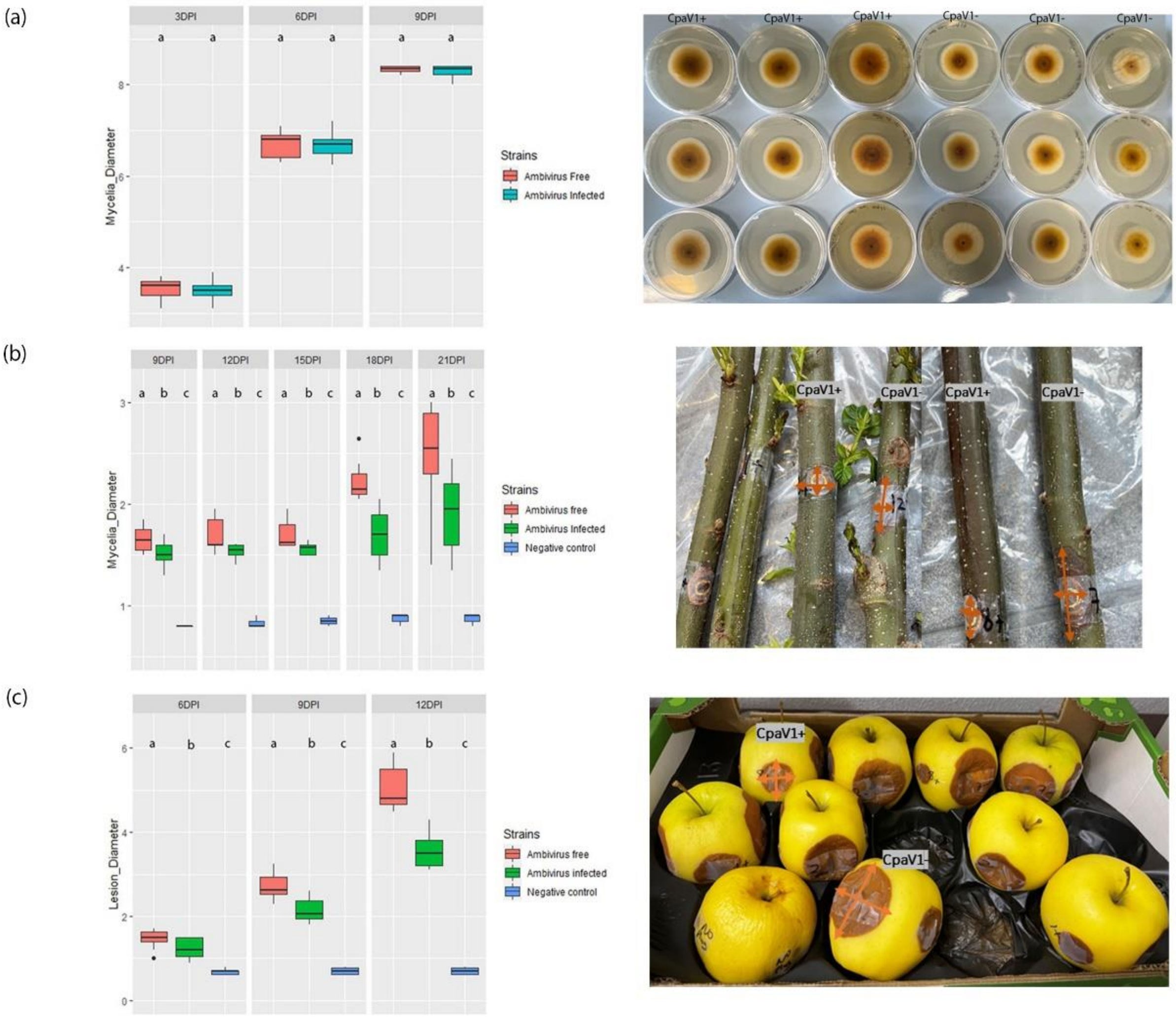


**Fig. S6. Cryphonectria parasitica ambivirus 1 (CpAV1) causes hypovirulence on chestnut cuttings and apples, but has no effect on phenotype and growth rate in axenic potato dextrose agar cultures.** (**A**) In vitro culture of ambivirus-infected and ambivirus-free *Cryphonectria parasitica* isolates in PDA (Sigma) at 7dpi; (**B**) Cankers caused by CpAV1-free and CpAV1-infected isolates at 9, 12, 15, 18 and 21 (picture) days post-inoculation. (**C**) Lesion diameters on apples at 6, 9 and 12 (picture) dpi; each apple displays two inoculated sites with a virus-positive isolate, and two inoculated sites with a virus-negative isolate. In (B) and (C), orange arrows represent diameters. The left side of each panel displays violin plots of measurements, the right-side exemplary photographs of the experiment. The virulence experiment on apples and the growth experiment on PDA was repeated twice, whereas the chestnut cuttings virulence experiment was carried out once; all the experiments were carried out with 9 biological replicates for each treatment (virus-infected, virus-free).

**Table S1.**

Ribozymes identified in ambivirus sequences available in GenBank.

| Ambivirus | Ribozyme (+)* | Ribozyme (-)* | Accession number |
|---|---|---|---|
| Armillaria ambi-virus 3 | HHRz (286-340) | HHRz (4518-4523;1-75) | MW423812.1; **MW423813.1;** MW423811.1 |
| Armillaria borealis ambi-like virus 1 | HHRz (527-578) | HPRz (61-139) | MW423804.1 § |
| Armillaria borealis ambi-like virus 1 | HHRz (526-577) | HPRz (60-138) | MW423805.1 § |
| Armillaria borealis ambi-like virus 2 | HHRz (284-342) | HHRz (4524-4529;1-75) | **MW423806.1**; MW423807.1;MW423808.1; MW4238010.1§ |
| Armillaria borealis ambi-like virus 2 | HHRz (284-342) | HHRz (4520-4525;1-75) | MW423809.1 § |
| Armillaria ectypa ambi-like virus 1 | HHRz (4952-4989**;1-19**) | HPRz (4466-4545) | BK014418.1 § |
| Armillaria luteobubalina ambi-like virus 1 | incomplete genome | incomplete genome | BK014419.1 § |
| Armillaria mellea ambi-like virus 1 | HPRz (2206-2310) | HPRz (2623-2548) | BK014420.1 § |
| Armillaria mellea ambi-like virus 2 | HHRz (45-99) | HHRz (4226-4303) | BK014421.1§ |
| Ceratobasidium ambivirus 1 | HHRz (2390-2449) | HPRz (2568-2657) | MN793993.1 |
| Cryphonectria parasitica ambivirus 1 | HHRz (4613-4623; 1-49) | HHRz (196-249) | MT354566** |
| Heterobasidion ambi-like virus 1 | HPRz (509-610) | HPRz (52-153) | MZ502384.1§ |
| Heterobasidion ambi-like virus 2 | incomplete genome | incomplete genome | MZ502385.1§ |
| Heterobasidion ambi-like virus 3 | HHRz (45-105) | HPRz (4725-4794) | MZ502386.1 § |

| | | | |
|---|---|---|---|
| Heterobasidion ambi-like virus 4 | HHRz (273-324) | HPRz (4886-4913; 1-35) | MZ502387.1§ |
| Phlebiopsis gigantea ambi-like virus 1 | HPRz (582-685) | HPRz (76-183) | MZ448624.1 § |
| Rhizoctonia solani ambivirus 1 | HHRz (2270-2327) | HHRz (2465-2548) | MT354567.1 |
| Rhizoctonia solani ambivirus 2 | HHRz (1470-1527) | HHRz (1601-1695) | MT354568.1 |
| Tulasnella ambivirus 1 | HHRz (4517-4578) | HHRz (1-19;4673-4736) | MN793991.1 |
| Tulasnella ambivirus 2 | likely incomplete genome | HHRz (121-209) | MN793992.1 |
| Tulasnella ambivirus 3 | HHRz (82-139) | HPRz (262-343) | MN793994 *** |
| Tulasnella ambivirus 4 | HHRz (4903-4924;1-38) | HPRz (157-247) | MN793995.1 |
| Tulasnella ambivirus 5 | HHRz( 4610-4632;1-34) | HPRz (191-269) | MN793996.1 |

*Numbers in brackets indicate the nt positions of the region spanning the ribozyme and refer to the virus sequence with the reported accession number. The (+) polarity is defined as the RNA strand coding for the polymerase (ORFA). Sequence available in GenBank as negative strand are marked by §. HPRzs are denoted in red.

** This sequence includes a terminal repeat of 243 nt, therefore the size of monomeric genomic RNA (excluding such a repeat) is 4623 nt

** *This sequence includes a terminal repeat of 249 nt, therefore the the size of monomeric genomic RNA (excluding such a repeat) is 4869 nt

**Table S2.** Examples of Mito-like viruses showing putative circular genomes (based on k-mer repeats) and paired-ambisense ribozymes (based on InfeRNAl searches).

| Entry | Name | Size (nt) circ | Ribozyme (+) | Ribozyme (-) |
|---|---|---|---|---|
| MW648461.1 | Grapevine-associated mitovirus 14 | 3210 | Twister | Hammerhead |
| MW648460.1 | Grapevine-associated mitovirus 13 | 2968 | Twister | Twister |
| MZ969057.1 | *Fusarium asiaticum* mitovirus 7 | 2992 | Varkud Satellite | Varkud Satellite |
| MZ969058.1 | *Fusarium asiaticum* mitovirus 8 | 3038 | Varkud Satellite | Varkud Satellite |
| SRR13754975 | NODE_4086_length_2968_cov_23.415553 | 2919 circRNA | Twister | Twister |
| SRR13754975 | NODE_3725_length_3056_cov_75.035584 | 3007 circRNA | Hammerhead | Twister |
| SRR13754975 | NODE_4217_length_2942_cov_119.153474 | 2893 circRNA | Twister | Hammerhead |
| SRR6943136 | NODE_404_length_3356_cov_47.364606 | 3307 circRNA | Twister | Twister |
| SRR6943136 | NODE_433_length_3356_cov_47.808407 | 3307 circRNA | Twister | Twister |
| SRR6031102 | NODE_2044_length_3642_cov_403.108714 | 3593 circRNA | Twister | Twister |
| SRR6765931 | NODE_243_length_3085_cov_22.068061 | 3036 circRNA | Twister | Twister |
| SRR10849745 | NODE_851_length_3256_cov_13.070688 | 3207 circRNA | Varkud Satellite | Varkud Satellite |
| SRR11565847 | NODE_733_length_2946_cov_27.405499 | 2897 circRNA | Hammerhead | Hammerhead |
| SRR11565844 | NODE_1557_length_2946_cov_62.985033 | 2897 circRNA | Hammerhead | Hammerhead |
| SRR12756234 | NODE_868_length_3013_cov_19.082993 | 2964 circRNA | Twister | Twister |
| SRR11076227 | NODE_2923_length_3393_cov_16.577711 | 3344 circRNA | Varkud Satellite | Varkud Satellite |
| SRR11541436 | NODE_36_length_3163_cov_14.808927 | 3114 circRNA | Varkud Satellite | Varkud Satellite |
| SRR11565434 | NODE_346_length_3332_cov_11.710341 | 3283 circRNA | Varkud Satellite | Varkud Satellite |
| SRR11565436 | NODE_673_length_3332_cov_331.066278 | 3283 circRNA | Varkud Satellite | Varkud Satellite |
| SRR12610547 | NODE_89_length_3256_cov_6.980522 | 3207 circRNA | Varkud Satellite | Varkud Satellite |
| SRR7142037 | NODE_1346_length_3408_cov_590.197601 | 3359 circRNA | Varkud Satellite | Varkud Satellite |

**Table S3.** Oligonucleotides used in this study and their features

| Primer name | Primer sequence (5 -3) | Primer position* | Virus | ID virus | Ribozyme | Used for | PCR amplicon (nt) |
|---|---|---|---|---|---|---|---|
| CryAMB1_HHp_5F | CCGTTCTATCAGCAAGTTCGT | 4525-4545 | Cryphonectria parasitica ambivirus 1 | MT354566.2 ** | HHRz (+) | PCR amplification of cDNA fragment containing the ribozyme | |
| | | | | | | | 299 |
| CryAMB1_HHp_6R | ATCGCTACTCCCTAATGCCC | 200-181 | Cryphonectria parasitica ambivirus 1 | MT354566.2 | HHRz (+) | PCR amplification of cDNA fragment containing ribozyme and 5'RACE of 3' RNA fragment generated during *in vitro* transcription | |
| CryAMB1_HHm_7F | CGTCGGGTGTGGCTGAAT | 98-115 | Cryphonectria parasitica ambivirus 1 | MT354566.2 | HHRz (-) | PCR amplification of cDNA fragment containing ribozyme and 5'RACE of 3' RNA fragment generated during *in vitro* transcription | |
| | | | | | | | 287 |
| CryAMB1_HHm_8R | GGTGAGGGTGAGGTTGATCTT | 384-364 | Cryphonectria parasitica ambivirus 1 | MT354566.2 | HHRz (-) | PCR amplification of cDNA fragment containing the ribozyme | |
| TuAV4_HH_9F | GCTTATGCCTGTTGTGCCTAC | 4792-4812 | Tulasnella ambivirus 4 | MN793995.1 | HHRz (+) | PCR amplification of cDNA fragment containing the ribozyme | |
| | | | | | | | 376 |
| TuAV4_HH_10R | GGTCCTACTGCATACACCGTA | 243-223 | Tulasnella ambivirus 4 | MN793995.1 | HHRz (+) | PCR amplification of cDNA fragment containing ribozyme and 5'RACE of 3' RNA fragment generated during *in vitro* transcription | |
| TuAV4_HP_11F | CTGTTTCGCATGTCCTTGGAA | 107-127 | Tulasnella ambivirus 4 | MN793995.1 | HPRz (-) | PCR amplification of cDNA fragment containing ribozyme and 5'RACE of 3' RNA fragment generated during *in vitro* transcription and 5' RACE of the viral RNA of (-) polarity strand | |
| | | | | | | | 319 |
| TuAV4_HP_12R | AGAAGAAGAAGACCAAGGCGA | 425-404 | Tulasnella ambivirus 4 | MN793995.1 | HPRz (-) | PCR amplification of cDNA fragment containing the ribozyme | |
| TuAV1_HHp_13F | AGAGAAGGTTATGTCGGGAGA | 4342-4362 | Tulasnella ambivirus 1 | MN793991.1 | HHRz (+) | PCR amplification of cDNA fragment containing the ribozyme | |
| | | | | | | | 347 |
| TuAV1_HHp_14R | CGAGACGAAACACACTTCTGG | 4688-4668 | Tulasnella ambivirus 1 | MN793991.1 | HHRz (+) | PCR amplification of cDNA fragment containing ribozyme and 5'RACE of 3' RNA fragment generated during *in vitro* transcription | |
| TuAV1_HHm_15F | CCATGTCGCACTGTCTGTATG | 4614-4594 | Tulasnella ambivirus 1 | MN793991.1 | HHRz (-) | PCR amplification of cDNA fragment containing ribozyme and 5'RACE of 3' RNA fragment generated during *in vitro* transcription | |
| | | | | | | | 388 |
| TuAV1_HHm_16R | TGTTGAGGAGATTCGTTGGGA | 245-225 | Tulasnella ambivirus 1 | MN793991.1 | HHRz (-) | PCR amplification of cDNA fragment containing the ribozyme | |

| | | | | | | |
|---|---|---|---|---|---|---|
| 30310-4488F | TGGTCCACCTTACACACCAG | 4500-4520 | Tulasnella ambivirus 1 | MN793991.1 | 5'RACE of the viral RNA of (-) polarity strand | |
| 30310-4341F | TGTCGGGAGATTCGTTTGAA | 4353-4272 | Tulasnella ambivirus 1 | MN793991.1 | 5'RACE of the viral RNA of (-) polarity strand | |
| 30310-320R | GAGTCGCAGTCGATCCGAAA | 351-332 | Tulasnella ambivirus 1 | MN793991.1 | 5'RACE of the viral RNA of (+) polarity strand | |
| 30310-289R | CGACAAGAACTTTTCGGGGC | 289-270 | Tulasnella ambivirus 1 | MN793991.1 | 5'RACE of the viral RNA of (+) polarity strand | |
| TuAV4_4624_For | GGCAGTGGACTTAGGCTTTAT | 4626-4644 | Tulasnella ambivirus 4 | MN793995.1 | 5'RACE of the viral RNA of (-) polarity strand | |
| TuAV4_321Rev | GAGGCTGATTGGTAGTGCAT | 321-302 | Tulasnella ambivirus 4 | MN793995.1 | 5'RACE of the viral RNA of (+) polarity strand | |
| TuAV4_248Rev | TGTGCGGTCCTACTGCATAC | 248-229 | Tulasnella ambivirus 4 | MN793995.1 | 5'RACE of the viral RNA of (+) polarity strand | |
| AmbiCp_RACE_4450_For | GTAGCACAAGTACCGAGTAGTC | 4571-4592 | Cryphonectria parasitica ambivirus 1 | MT354566.2 | 5'RACE of the viral RNA of (-) polarity strand | |
| AmbiCp_RACE_4403_For | TTGTCGTTAAGGACATACCTGGGT | 4545-4568 | Cryphonectria parasitica ambivirus 1 | MT354566.2 | 5'RACE of the viral RNA of (-) polarity strand | |
| AmbiCp_RACE_267_Rev | GAGAGCTCGTCGTGAAGAG | 409-391 | Cryphonectria parasitica ambivirus 1 | MT354566.2 | 5'RACE of the viral RNA of (+) polarity strand | |
| AmbiCp_RACE_229_Rev | TTGATCTTGATGACCTCCTGGC | 371-350 | Cryphonectria parasitica ambivirus 1 | MT354566.2 | 5'RACE of the viral RNA of (+) polarity strand | |
| DN30310-real-F | TGGGGACCAACAGATGAACG | 2644-2663 | Tulasnella ambivirus 1 | MN793991.1 | Stranded qPCR | |
| | | | | | | 107 |
| DN30310-Real-Rev | GGGGCAAAACGACACAAGTC | 2751-2732 | Tulasnella ambivirus 1 | MN793991.1 | Stranded qPCR | |
| 30310-real136F | GGGGGTCTAACCAGAAGTGTG | 4658-4678 | Tulasnella ambivirus 1 | MN793991.1 | Stranded qPCR | |
| | | | | | | 113 |
| 30310-real230R | GCATGATGAATGGTCCAGTG | 35-16 | Tulasnella ambivirus 1 | MN793991.1 | Stranded qPCR | |
| 30310-real4795R | CGACATGGGGTAGTCACATTT | 4601-4581 | Tulasnella ambivirus 1 | MN793991.1 | Stranded qPCR | 101 |
| 30310-4488F | TGGTCCACCTTACACACCAG | 4500-4520 | Tulasnella ambivirus 1 | MN793991.1 | 5'RACE of the viral RNA of (-) polarity strand and stranded qPCR | |

* 5' -3' nt positions refereed to the genomic viral sequence in GenBank (ID virus)

** in MT354566.2 the region from 4624 to 4866 is a direct repetition of positions 1 to 243, therefore the end of the full-length genome is from positions 1 to 4623
